# Modification of oxygen consumption and blood flow in mouse somatosensory cortex by cell-type-specific neuronal activity

**DOI:** 10.1101/651224

**Authors:** Matilda Dahlqvist, Kirsten Thomsen, Dmitry Postnov, Martin Lauritzen

**Affiliations:** Department of Neuroscience, University of Copenhagen, Blegdamsvej 3, DK-2200 Copenhagen N, Denmark; Department of Neurophysiology, Rigshospitalet Glostrup, Valdemar Hansens Vej 1-23 DK-2620 Glostrup, Denmark; Department of Biomedical Sciences, University of Copenhagen, Blegdamsvej 3, DK-2200 Copenhagen N Denmark

## Abstract

Gamma activity arises from the interplay between pyramidal neurons and fast-spiking parvalbumin (PV) interneurons, is an integral part of higher cognitive functions and is assumed to contribute importantly to brain metabolic responses. Cerebral metabolic rate of oxygen (CMRO_2_) responses were evoked by optogenetic stimulation of cortical PV interneurons and pyramidal neurons. We found that CMRO_2_ responses depended on neuronal activation, but not on the power of gamma activity induced by optogenetic stimulation. This implies that evoked gamma activity *per se* is not energy demanding. Optogenetic stimulation of PV interneurons during somatosensory stimulation reduced excitatory neuronal activity but did not potentiate O_2_ consumption as previously hypothesized. In conclusion, our data suggest that activity-driven CMRO_2_ responses depend on neuronal excitation rather than the cerebral rhythmic activity they induce. Excitation of both excitatory and inhibitory neurons requires energy, but inhibition of cortical excitatory neurons by interneurons does not potentiate activity-driven energy consumption.

## Introduction

FS PV-expressing interneurons are associated with the formation of network gamma oscillations (30- 90 Hz) ^1–3^, which are fundamental for cognition, perception and memory formation ^4–6^. Gamma activity results from the interaction between pyramidal cells and FS PV interneurons, where PV interneuron synchronization creates ‘windows of opportunity’ for pyramidal neuron spiking ^1,2,7^.

Previous studies have indicated a strong link between the BOLD signal and the gamma component of the local field potential (LFP) band ^8,9^. In addition, more recent studies have described a link between gamma oscillations and the oxygenation of brain tissue where ultra-slow fluctuations enveloping the gamma band modulate vasomotion ^10,11^. Since PV interneurons are fast spiking, they have a greater energy demand than regular spiking neurons ^12^ and are dependent upon a continuous delivery of oxygen and glucose to provide sufficient amounts of ATP ^13,14^. It is thought that deficits of either oxygen or glucose supply to PV interneurons will decrease gamma activity ^15^.

Assessing oxygen consumption due to interneuron inhibition in intact neuronal networks is complicated by non-linear gain modulation of principal cell excitatory input, resulting in progressively increasing feedforward inhibition of PV interneurons with progressively increasing excitatory drive ^16^. Increasing near-concomitant excitation and hyperpolarizing inhibition potentiates ion displacement across the cell membrane without necessarily affecting either spike rate or membrane potential ^17^. Due to the potentiated ion displacement, increasing excitation and inhibition concomitantly is hypothesized to augment oxygen consumption non-linearly ^17^.

PV interneurons may also play a role in generating stimulus-evoked CBF responses, as optogenetic stimulation of GABAergic interneurons induces CBF responses that are not due to disinhibition of surrounding pyramidal neurons ^18–21^. While some interneurons express vasoactive substances thought to contribute to stimulation-induced blood flow responses ^19,22^, no vasoactive substances are known to be released by PV interneurons. Nonetheless, some studies have shown that PV interneurons do influence blood flow and artery diameter ^23,24^.

Taken together, PV interneurons appear to be centrally placed between gamma activity, brain oxygen consumption and cerebral blood flow. In the present study, we examined the relationship between PV interneuron activity, gamma activity and brain oxygen use *in vivo*. Due to the high energy demands of PV interneurons generating gamma oscillations, we hypothesized that stimulation-induced network gamma oscillations determine CMRO_2_. We found that *in vivo* optogenetic stimulation of PV interneurons expressing channelrhodopsin-2 (ChR2) did induce CMRO_2_ responses and did so independently of pyramidal neuron or other interneuron involvement. Whisker pad stimulation of pyramidal neurons induced gamma activity that was halved by concurrent optogenetic stimulation of PV interneurons, while CMRO_2_ responses remained unchanged. Thus, gamma oscillations *per se* did not evoke greater oxygen consumption than the activity of the constituent neurons of the gamma circuit. We also examined the notion that concurrent excitation and inhibition evoke a larger ion flux than the sum of ion fluxes evoked by excitation and inhibition separately, which is the basis of the non-linear gain modulation of principal cell excitation. We found that the excitation-inhibition balance of the neural network was an important determinant of CMRO_2_ response amplitude to whisker pad stimulation and that CMRO_2_ responses could not be used as an indirect measure of stimulus-evoked ion flux.

## Results

Whisker pad (WP) stimulation involves a multisynaptic signalling pathway that initially targets L4 pyramidal neurons of the barrel cortex via the ventral posterior thalamic nucleus, and subsequently projects to L2/3 and L5 ^25,26^. Thalamocortical as well as inter- and intracortical input to the barrel cortex is glutamatergic and is modulated by interneuronal GABAergic activity in the form of disinhibition, feedforward and feedback inhibition ^16,27^. Optogenetic stimulation bypasses the initial part of the somatosensory signalling pathway leading to L4, L2/3 or L5 target cells and therefore interrogates post-target cell processing. In the present study, we have employed optogenetic activation of channelrhodopsin (ChR2)-expressing pyramidal neurons and PV interneurons as well as WP stimulation to investigate gamma generation and the relation between gamma activity, neuron and interneuron activation, and cortical oxygen use. Simultaneous activation of pyramidal neurons by WP stimulation and optogenetic activation of PV interneurons was employed to evaluate the effect of inhibitory input on excitation. CMRO_2_ responses were taken as expression of the workload necessary to restore ionic gradients after depolarisation.

### Characterization of cells containing channelrhodopsin

To directly activate neurons and interneurons that are part of the sensory input processing pathway, we employed two mouse lines, one with ChR2-expressing PV interneurons (PV/ChR2), the other with ChR2-expressing pyramidal neurons (Pyramidal/ChR2). In both mouse strains, ChR2 was tethered to enhanced yellow fluorescent protein (EYFP). Twenty μm thick coronal slices from layer L2/3 of the whisker barrel cortex (bregma +0.5 mm/-1.94 mm ^28^) were examined. In PV/ChR2 brains, co-localisation of parvalbumin with EYFP was seen in > 90% of PV interneuron somata and processes (fig. S1d-f). EYFP rings depicting cells expressing ChR2 (fig. S1f) were only seen in connection with PV interneurons. In Pyramidal/ChR2 brain, EYFP expression was intensive due to labelling of the ubiquitous processes of pyramidal neurons and EYFP rings were not distinguishable (fig. S1b). Pyramidal neuron somata were found in all neocortical layers from L2/3 to L5 (fig. S1a showing L2/3).

### Optogenetic stimulation of PV interneurons or pyramidal neurons under control conditions induces O_2_ use and increases local blood flow

The whisker barrel cortex was activated in three ways, by **1)** direct optogenetic activation of either PV interneurons or pyramidal neurons (Optogenetic PV or Optogenetic Pyramidal stimulation, fig 1c, f), **2)** somatosensory stimulation of the contralateral whisker pad (WP_PV_ or WP_PYR_ stimulation, fig 1e) or **3)** combined direct optogenetic activation and concurrent WP stimulation targeting the whisker barrel cortex of the same hemisphere (Combined PV or Pyramidal stimulation, fig 1d, g). Note that in the case of Combined PV stimulation, both PV interneurons and pyramidal neurons were activated, while in the case of Combined Pyramidal stimulation, only pyramidal neurons were activated (see Materials and Methods).

**Figure 1:**
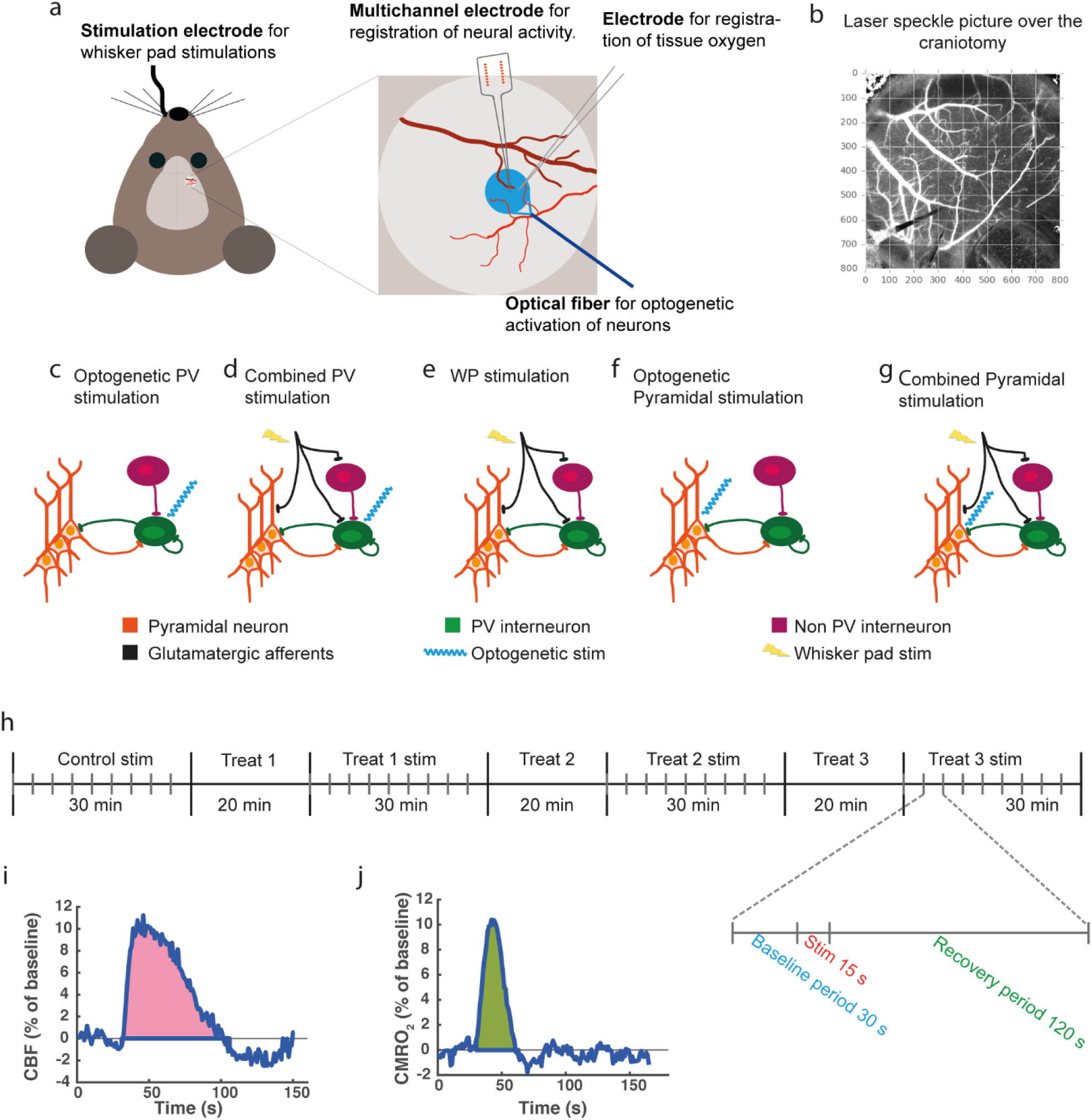
Experimental setup. (a) A schematic of the experimental setup showing placement of multichannel electrode measuring extracellular electrical activity, oxygen electrode measuring tissue oxygen partial pressure, optical fibre used to activate ChR2-expressing neurons with blue light (473 nm) and electrode in the whisker pad for infraorbital nerve stimulation. An optical fiber with red laser light (785 nm) and laser speckle camera used for measuring blood flow were positioned ~9 cm above the craniotomy and are therefore not included in the schematic. The blue circle indicates the cortical area stimulated during optogenetic illumination. (b) A laser speckle image of the craniotomy, where placement of the measuring electrodes and the optical fibre can be discerned in the lower left corner. (c-g) Schematics showing the stimulation types used in the present study. (c) Optogenetic PV stimulation: blue laser light directly activates PV interneurons. (d) Combined PV stimulation: WP stimulation excites pyramidal neurons and PV interneurons via thalamocortical afferents while PV interneurons are additionally directly activated by blue laser light. (e) WP stimulation excites pyramidal neurons and interneurons via thalamocortical afferents. (f) Optogenetic Pyramidal stimulation: blue laser light directly activates pyramidal neurons. (g) Combined Pyramidal stimulation: WP stimulation activates pyramidal neurons and PV interneurons via thalamocortical afferents while pyramidal neurons are additionally directly activated by blue laser light. (h) A schematic of the experimental protocol showing the 4 stimulation periods interleaved with three periods in which treatment was applied and allowed to work. There was no wash-out between treatments. Thus, treatments were cumulative. The individual stimulation protocol is also shown with 30 s baseline before and 120 s recovery period after each 15 s stimulation train. (i-j) Typical CBF and CMRO_2_ responses showing how AUC_CBF_ (i) and AUC_CMRO2_ (j) were calculated.

In control conditions, we found that stimulation of PV interneurons or pyramidal neurons using any of the three stimulation types evoked robust metabolic (fig. 2c, d) and neurovascular responses (fig. 2a, b). Optogenetic activation of PV interneurons also diminished the sparse spontaneous gamma activity present at baseline (fig 2e) as well as the gamma activity evoked by whisker pad stimulation during Combined PV stimulation (compare WP_PV_ vs Combined PV stimulation, fig 2e), confirming that the output of PV interneurons induces inhibition. In the same mice, Combined PV stimulation and WP_PV_ stimulation evoked CMRO_2_ responses that were of equal magnitude, although Combined PV stimulation was characterized by greater inhibition due to the optogenetic activation of PV interneurons (fig 2c). Combined PV stimulation also evoked LFP amplitudes that were half as large as those induced by WP_PV_ stimulation (fig. 3a, c), indicating that Combined PV stimulation induced less net excitation of pyramidal neurons compared to WP_PV_ stimulation.

**Figure 2.**
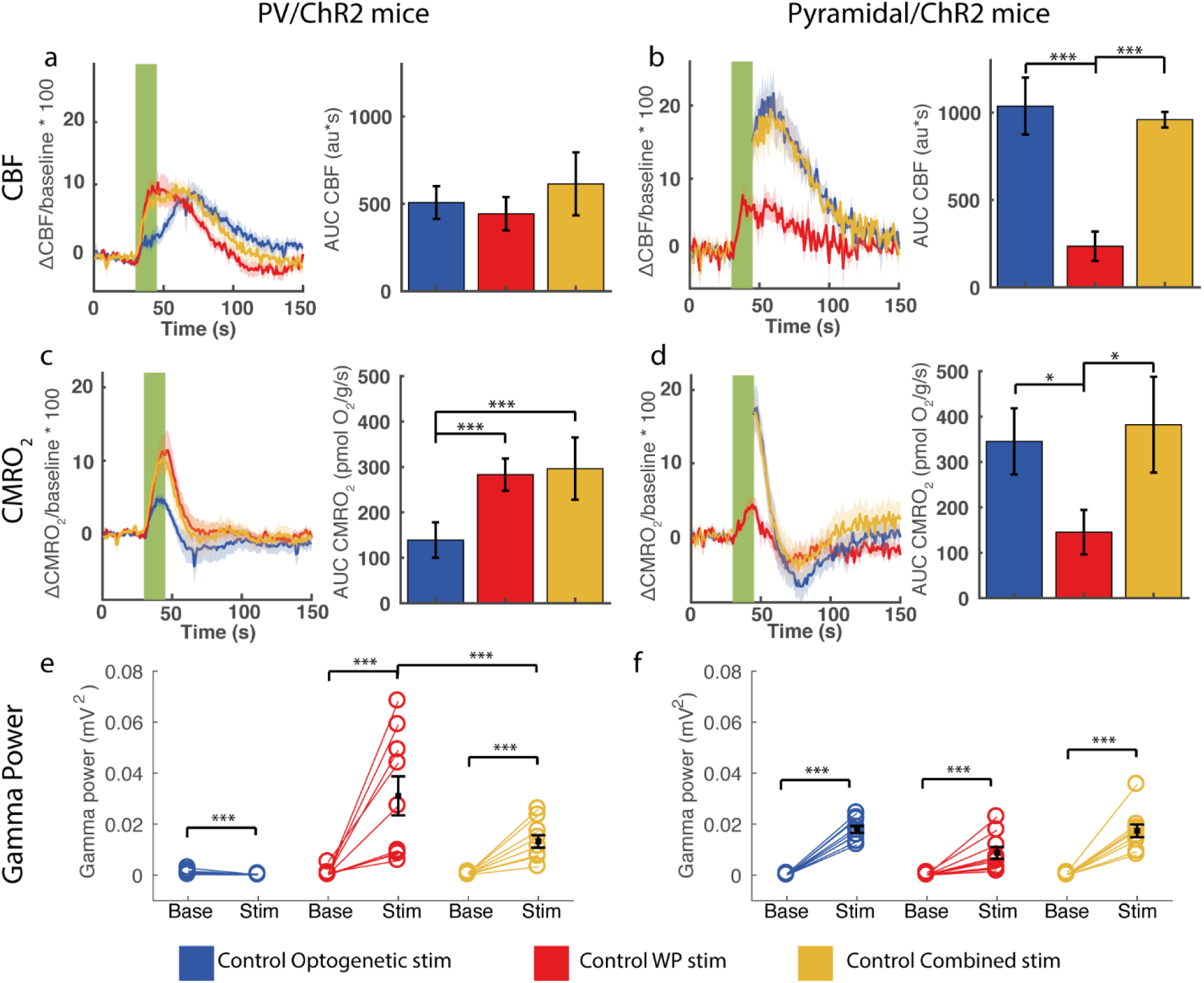
Evoked CBF, CMRO_2_ and gamma frequency activity during control conditions in PV/ChR2 mice and Pyramidal/ChR2amidal mice. Evoked CBF responses in (a) PV/ChR2 mice and (b) Pyramidal/ChR2 mice. Left panels (a, b) show CBF time courses during optogenetic, WP and Combined stimulations and right graphs (a, b) show the corresponding calculated AUC_CBF_. Note the biphasic CBF response to optogenetic stimulation in PV/ChR2 mice, which peaks ~21 s after the CBF responses to WP and Combined PV stimulations. Evoked CMRO_2_ responses in (c) PV/ChR2 mice and (d) Pyramidal/ChR2 mice. Left panels (c, d) show CMRO_2_ time courses during optogenetic, WP and Combined stimulations and right panels (c, d) show the corresponding calculated AUC_CMRO2_. Evoked gamma activity in (e) PV/ChR2 mice and (f) Pyramidal/ChR2 mice during baseline (to the left) and stimulation (to the right). The CBF and CMRO_2_ stimulation time courses are averaged across animals. Each CBF time course was extracted from a roi (40 pixels in diameter) placed next to the insertion points of the oxygen electrode and the multichannel electrode in the laser speckle recordings and used for calculating CMRO_2_ time courses, AUC_CBF_ and AUC_CMRO2_. Green fields represent 15 s stimulation trains. We were not able to register the CBF signal during optogenetic stimulation in Pyramidal/ChR2 mice. All data is presented as mean ± SEM. n = 9 for all groups. Significant difference is shown by asterisk(s): *P < 0.05, **P < 0.01 and ***P < 0.005.

**Figure 3:**
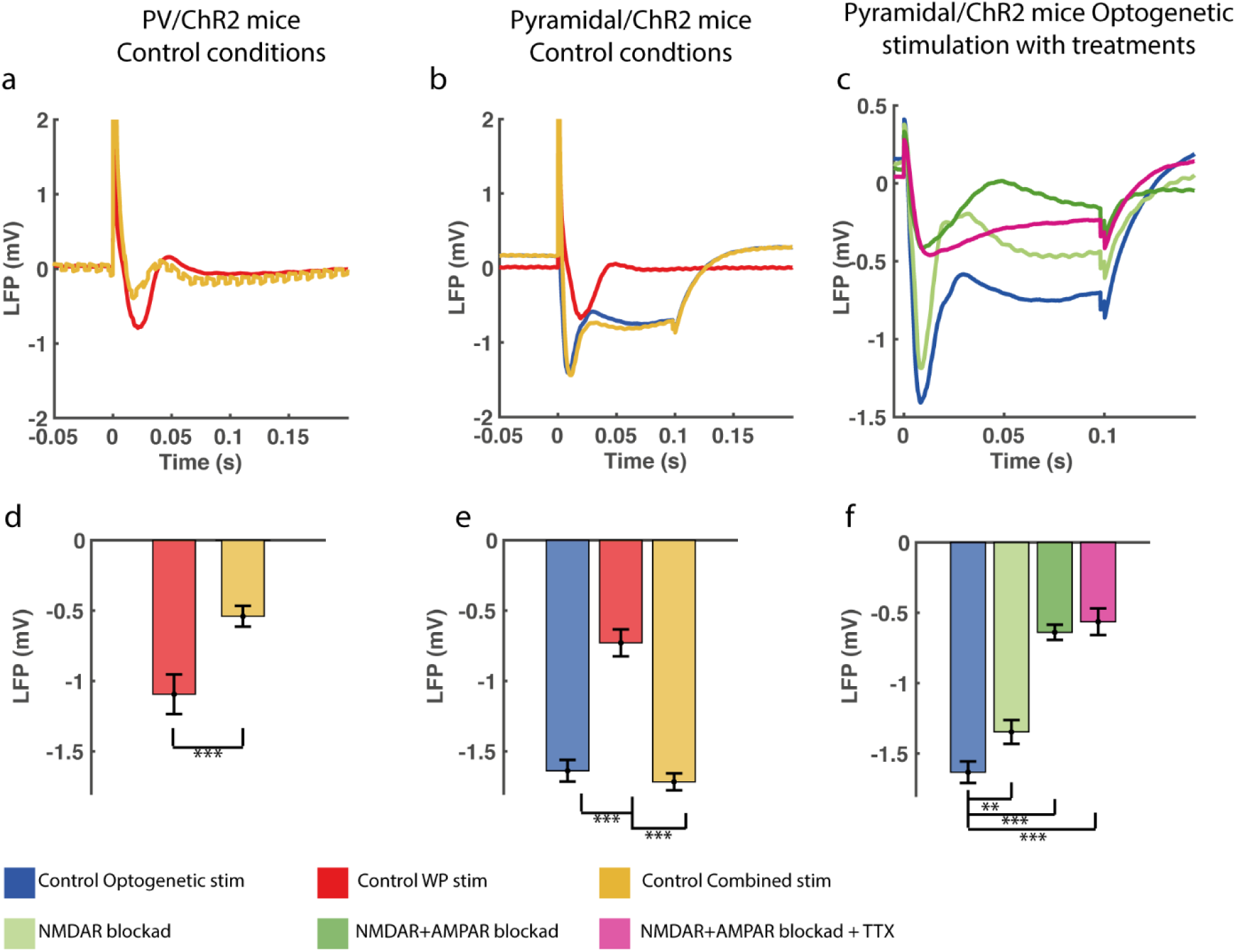
Local field potentials during control conditions in PV/ChR2 and Pyramidal/ChR2 mice and during all treatments in Pyramidal/ChR2 mice. (a, d) In PV/ChR2 mice in control conditions, LFP amplitude evoked by WP stimulation was significantly reduced by simultaneous optogenetic stimulation of PV interneurons (Combined PV stimulation), demonstrating that optogenetic stimulation activated PV interneurons and increased inhibitory input to pyramidal neurons. Combined PV stimulation gave a jagged LFP trace due to the optogenetic stimulation frequency of 100 Hz. (b, e) In Pyramidal/ChR2 mice in control conditions, optogenetic stimulation of pyramidal neurons (impulse duration, 0.1 s) evoked a rapid deflection which partially decayed during illumination followed by a slight rebound hyperpolarization after termination of the stimulus. The same response pattern was seen during Combined Pyramidal stimulation. WP stimulation-evoked LFPs from the same mice is also shown. WP stimulation-evoked LFP amplitudes from the two mouse strains were not significantly different. (c, f) In Pyramidal/ChR2 mice, blocking NMDAR reduced LFP amplitude and greatly so after the subsequent application of AMPAR blocker. LFP amplitude was not altered by the further application of TTX. We surmise that LFP in the presence of TTX represents the opening and gradual closing of ChR2 pores.

Optogenetic activation of pyramidal neurons and Combined Pyramidal stimulation induced substantial CMRO_2_ responses. In contrast, CMRO_2_ responses to WP_Pyr_ stimulation were smaller (WP_Pyr_ vs Optogenetic Pyramidal: p=0.048, WP_Pyr_ vs Combined Pyramidal: p=0.014), although all three stimulation types targeted pyramidal neurons of the same whisker barrel cortex. This finding may reflect either greater numbers of activated neurons or greater ion fluxes due to the opening of ChR2 channels in activated neurons during optogenetic pyramidal neuron activation and Combined Pyramidal stimulation. This is indicated by the greater evoked gamma activities and LFPs during Optogenetic Pyramidal and Combined Pyramidal stimulations than during WP_Pyr_ stimulation (fig 2 f+3b, d).

Thus, we found that under control conditions, optogenetic activation of PV interneurons evoked substantial CMRO_2_ and CBF responses without evoking gamma activity; that evoked CMRO_2_ responses and gamma activity were proportionate during optogenetic activation or whisker pad stimulation of pyramidal neurons alone; and that activation of pyramidal neurons and PV interneurons together disrupted this relation evoking less gamma activity without diminishing CMRO_2_ responses.

Despite differences in CMRO_2_ responses, evoked LFP amplitudes and gamma activity, Optogenetic PV stimulation, WP_PV_ stimulation and Combined PV stimulation all evoked CBF responses of the same amplitude and AUC_CBF_ (fig 2a, fig. S2a). Notably, CBF responses evoked by Combined PV stimulation were not less than CBF responses evoked by WP_PV_ stimulation alone despite decreased LFP amplitude and gamma activity (described above), implying that mechanisms distinct from postsynaptic excitation determine CBF response magnitude. Also, the time course of CBF responses differed between Optogenetic PV and WP_PV_ stimulations. Optogenetic PV stimulation evoked a biphasic CBF response (fig. 2a), which reached its maximum 21.0 ± 3.4 s later than the monophasic CBF responses evoked by WP_PV_ stimulation (fig. S2b). In contrast, the CBF responses in Pyramidal/ChR2 mice to all stimulation types in control conditions paralleled the CMRO_2_ responses (fig 2b). Since there was no difference between Optogenetic Pyramidal and Combined Pyramidal responses, we will from now on only refer to Optogenetic Pyramidal stimulation results. WP stimulation evoked unequal CBF responses in the two mouse strains, (WP_PV_ vs WP_PYR_, p = 0.039; fig. 2 a, b), although the WP stim-evoked LFPs in the two strains were statistically not different (p = 0.139; fig. 3c, d). Thus, although the excitatory input to pyramidal neurons in PV/ChR2 and Pyramidal/ChR2 mice was similar, post-synaptic processing of the signal leading to vasodilation in the two mouse strains differed.

### Blockade of NMDAR, AMPAR and GABA_A_R potentiated O_2_ use while blockade of voltage-gated Na^+^ channels abolished it in PV/ChR2 mice

To ensure that the responses described above represent innate properties of PV interneurons and not neuronal interactions, we applied iGluR and GABA_A_R blockers (MK801+NBQX and GABAzine, respectively) topically to the brain to inhibit collateral stimulation, inhibition and disinhibition. Lastly, we applied tetrodotoxin (TTX) to block voltage-gated Na^+^ channels. We found that the CMRO_2_ and CBF responses to Optogenetic PV stimulation were not by-products of interneuron or pyramidal neuron interactions as these responses were also present during iGluR+GABA_A_R blockade (fig. 4a, b). Thus, PV interneuron activity *per se* was found to be energy demanding and able to induce CBF responses.

**Figure 4.**
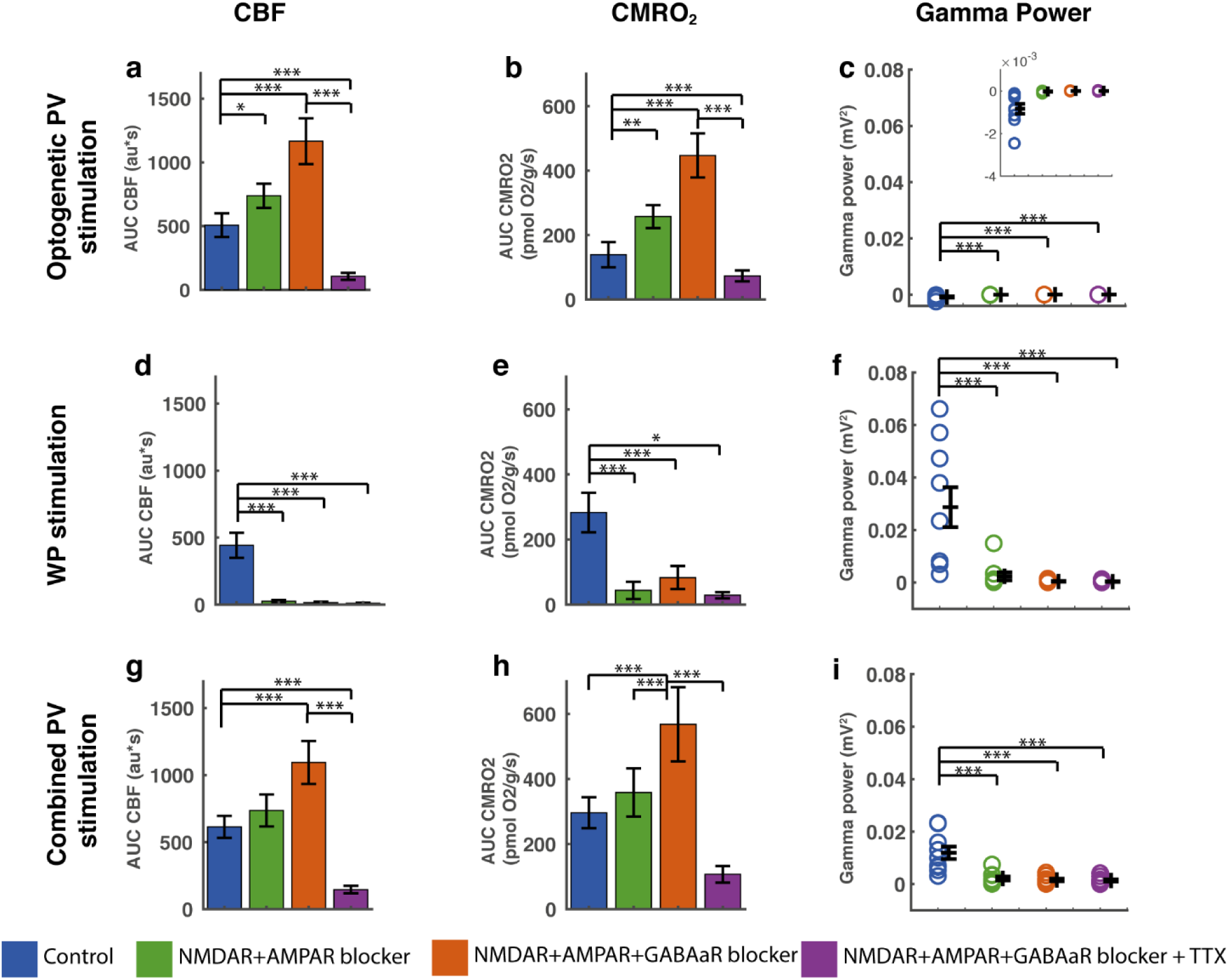
Evoked CMRO_2_, CBF and gamma power responses during blockade of NMDAR, AMPAR, GABA_A_R and Na^+^ channels in PV/ChR2 mice. Optogenetic PV stimulation-evoked CBF, CMRO_2_ and gamma power responses in (a - c), WP stimulation-evoked CBF, CMRO_2_ and gamma power responses in (d - f) and Combined PV stimulation-evoked CBF, CMRO_2_ and gamma power responses in (g - i). CBF and CMRO_2_ responses are calculated as AUC. The evoked gamma power responses shown are calculated as the gamma power responses minus baseline gamma values. Ionotropic GluR blockade abolished evoked gamma power for all stimulation types (c, f, i) and abolished CBF and CMRO_2_ responses to WP stimulation (d, e). Ionotropic GluR+GABA_A_R blockers augmented and subsequent TTX abolished both CBF and CMRO_2_ responses to Optogenetic PV and Combined PV stimulation. Data is presented as mean ± SEM. n = 9 for all groups. Significant differences are shown by asterisk(s): *P < 0.05, **P < 0.01 and ***P < 0.005.

In PV/ChR2 mice, iGluR blockade significantly increased CMRO_2_ and CBF responses to Optogenetic PV stimulation (fig. 4a, b). In contrast, the CMRO_2_ and CBF responses to Combined PV stimulation during iGluR blockade were not significantly potentiated (fig. 4g, h). Ionotropic GluR blockers abolished all responses to WP_PV_ stim indicating that NMDAR and AMPAR activation was indeed suppressed (fig. 4d, e; ^29^. Abolishing the GABA_A_ergic inhibitory tonus further potentiated both CMRO_2_ and CBF responses, as this inhibitory tonus affects all cortical neuron types. Lastly, application of TTX nearly abolished CMRO_2_ and CBF responses to Optogenetic PV and Combined PV stimulations (fig 4a, b, g, h), indicating the almost 100% dependence of these responses on voltage-gated Na^+^ channels.

### Blockade of NMDAR and AMPAR increases the evoked CBF while the evoked O_2_ use is unchanged in Pyramidal/ChR2 mice

A slightly different treatment protocol was followed in Pyramidal/ChR2 mice. Initially, NMDAR blockade alone was applied to evaluate the relation between NMDAR and gamma activities. Following this, AMPAR blocker was added to achieve the same iGluR blockade as in PV/ChR2 mice and lastly, TTX was applied. We found that blocking NMDAR alone did not significantly affect CMRO_2_ or CBF responses to either optogenetic or WP pyramidal neuron activation, but did reduce LFP amplitude by 17 and 26% (fig. 3f; LFP_WP_ not shown) and gamma activity by 23 and 37% (see below; fig. 5 a, b, c, d, e, f). Ionotropic GluR blockade abolished CBF or CMRO_2_ responses to WP stimulation in Pyramidal/ChR2 mice as expected (fig. 5d, e). In contrast, it potentiated the CBF response to Optogenetic Pyramidal neuron activation, but did not affect the corresponding CMRO_2_ response (fig 5a, b). Ionotropic GluR blockade reduced the evoked LFP amplitude to Optogenetic Pyramidal neuron activation and WP stimulation by 60% and 67%, but further application of TTX did not reduce LFP amplitude significantly (fig. 3f; LFP_WP_ not shown). TTX only slightly reduced the CBF responses to Optogenetic Pyramidal stimulation (fig. 5a), but reduced the corresponding CMRO_2_ response to Optogenetic Pyramidal stimulation by two-thirds (fig. 5b). Thus, a mis-match between CBF and CMRO_2_ responses to Optogenetic Pyramidal stimulation was found in Pyramidal/ChR2 mice. The small TTX-induced reduction in CBF suggests a minor dependency of these neurons on voltage-gated Na^+^ channels and that a large part of pyramidal neuron oxygen consumption is devoted to restoring ionic gradients after ChR2 channel opening.

**Figure 5.**
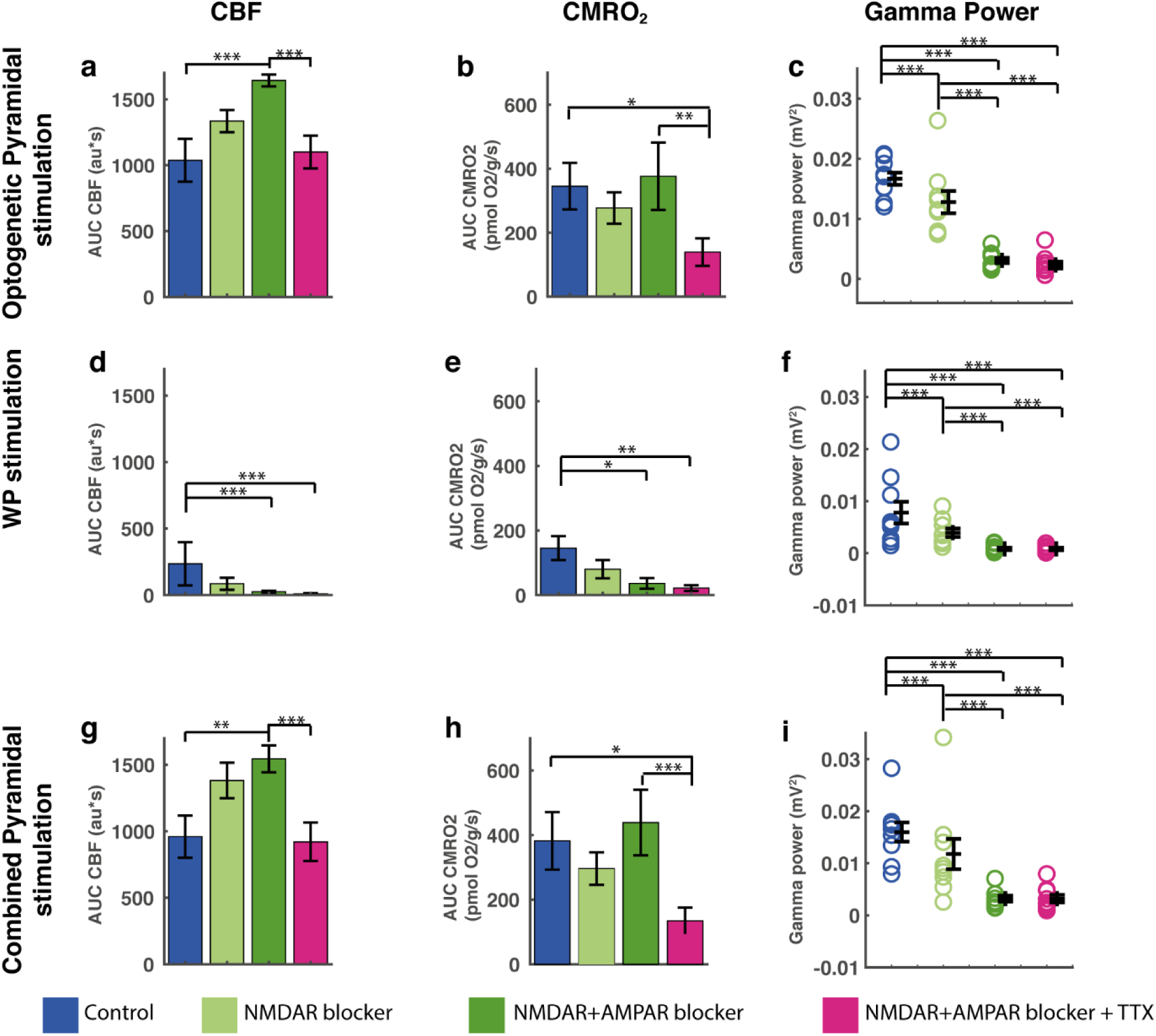
Evoked CMRO_2_, CBF and gamma power responses during blockade of NMDAR, AMPAR and Na^+^ channels in Pyramidal/ChR2 mice. Optogenetic Pyramidal stimulation-evoked CBF, CMRO_2_ and gamma power responses in (a - c), WP stimulation-evoked CBF, CMRO_2_ and gamma power responses in (d - f) and Combined Pyramidal stimulation-evoked CBF, CMRO_2_ and gamma power responses in (g - i). CBF and CMRO_2_ responses are calculated as AUC. The evoked gamma power responses shown are calculated as the gamma power responses minus baseline gamma values. Treatment with the NMDAR blocker MK801 decreased evoked gamma responses for all three stimulation types (c, f, i). Ionotropic GluR blockade increased evoked CBF responses to Optogenetic (a) and Combined (g) Pyramidal stimulation, while CMRO_2_ remained unchanged compared to controls (b, h). All evoked gamma responses were abolished by iGluR blockade (c, f, i). TTX reduced CMRO_2_ responses to Optogenetic and Combined Pyramidal stimulation (b, h), while CBF responses were unchanged compared to control values (a, g). Data is presented as mean ± SEM. n = 9 for all groups. Significant differences are shown by asterisk(s): *P < 0.05, **P < 0.01 and ***P < 0.005.

### Gamma activity was abolished with NMDAR and AMPAR blockade

We wondered if this difference in CBF and CMRO_2_ responses between the two mouse strains would be reflected in gamma activity. In both mouse strains, substantial gamma activity was only found when somatosensory or Optogenetic Pyramidal stimulation was part of the stimulation protocol ^30^(fig. 2, e, f; 4, b-f). In PV/ChR2 mice, optogenetic activation of PV interneurons during control circumstances reduced spontaneous, i.e. baseline, gamma activity as well as evoked gamma activity during WP_PV_ stimulation (fig. 4 c, f, i), confirming that optogenetic activation of PV interneurons induces inhibition of surrounding pyramidal neurons. In Pyramidal/ChR2 mice, all three stimulation types evoked gamma responses (fig. 5, c, f, i), the smallest response being evoked by WP_Pyr_ stimulation, corresponding to the smaller CBF and CMRO_2_ responses in these mice. A slight but significant decrease in gamma activity was seen for all three stimulation types after NMDAR blockade in these mice (decrease of ~29% when pooling all stimulation types, p = 4.69×10^-5^; fig. 5, c, g, i). For both mouse strains and all stimulation types, combined NMDAR and AMPAR blockade abolished gamma responses entirely. These results demonstrate the necessity of pyramidal neuron activation for the induction of gamma activity.

### Dissociation between oxygen consumption and gamma activity during optogenetic stimulation of PV interneurons and pyramidal neurons

To evaluate energy consumption of gamma activity, we plotted evoked CMRO_2_ responses as a function of gamma power (fig 6). In Pyramidal/ChR2 mice, a linear relationship across treatments was found between WP_Pyr_ stimulation-evoked CMRO_2_ responses on the one hand and corresponding gamma power responses on the other (fig. 6a). The same proportionality between CMRO_2_ responses and gamma activity was found for Optogenetic Pyramidal stimulation during control conditions. During NMDAR blockade, this relationship was not affected as CMRO_2_ responses and gamma activity decreased proportionally, but during optogenetic stimulation in the presence of AMPAR blockade, gamma activity diminished greatly with no reduction in CMRO_2_. No relationship was found between optogenetically evoked CMRO_2_ responses and gamma responses in PV/ChR2 mice. Specifically, in PV/ChR2 mice, large CMRO_2_ responses were elicited by Optogenetic PV stimulation in the presence of iGluR+GABAergic blockers without evoking any gamma responses (fig. 6b). Thus, in both mouse strains, iGluR±GABAergic blockers revealed a dissociation between oxygen consumption and gamma activity.

**Figure 6.**
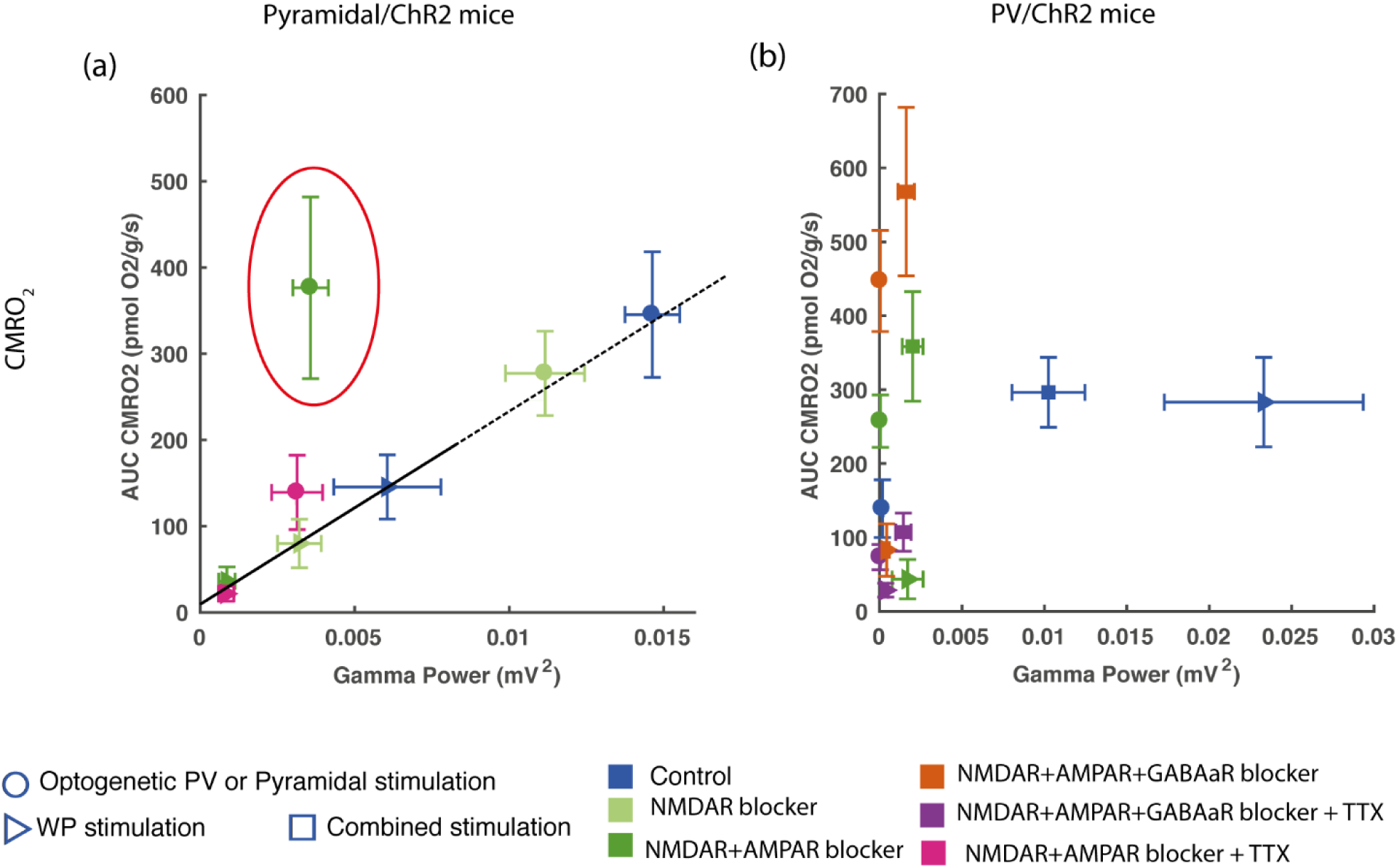
CMRO_2_ as a function of Gamma Power. (a) In Pyramidal/ChR2 mice, a linear relation between CMRO_2_ and gamma power was seen during control conditions and NMDAR blockade, implicating NMDAR activity as a modulator of gamma power. AMPAR blockade disrupted this relation as gamma power was nearly abolished without altering CMRO_2_ responses (AMPAR response encircled). (b) In PV/ChR2 mice, WP and Combined PV stimulation produced equal CMRO_2_ responses although the simultaneous stimulation of pyramidal neurons and PV interneurons during Combined PV stimulation inhibited gamma activity by more than 50%. No relationship between CMRO_2_ and gamma power in PV/ChR2 mice was evident.

## Discussion

While several optogenetic studies have looked at CBF responses to interneuronal activation ^18,19^, only few studies have studied the corresponding CMRO_2_ responses ^21^. These previous studies have employed a mouse strain in which ChR2 was expressed in all interneurons. In the present study, ChR2 expression was restricted to either PV interneurons or pyramidal neurons using two separate mouse strains. We report here that in control conditions, optogenetic stimulation targeting PV interneurons alone resulted in substantial CMRO_2_ as well as CBF responses. Notably, blocking iGluRs and GABA_A_Rs did not diminish response magnitude. On the contrary, in the presence of these blockers, optogenetic stimulation of PV interneurons evoked CMRO_2_ and CBF responses that were equally augmented compared to control conditions. In comparison, augmented CBF responses are not seen in the presence of iGluR and GABA_A_R blockers when all interneuron types are activated by optogenetic stimulation ^18,21^, in keeping with a stimulation-driven inhibitory action of non-PV interneurons on PV interneurons. The CMRO_2_ and CBF responses evoked by optogenetic stimulation of PV interneurons in the presence of iGluR and GABA_A_R blockers were almost abolished by subsequent application of TTX, indicating that these responses were dependent upon voltage-gated Na^+^ channels in agreement with the supercritical density of these channels in PV interneurons (Hu et Jonas, 2014).

CBF responses evoked by optogenetic stimulation of pyramidal neurons were also augmented by iGluR blockers. In contrast, application of TTX in the presence of iGluR blockers only slightly reduced the CBF response, suggesting that most of the CBF response to optogenetic stimulation of pyramidal neurons was due to cation flux through opened ChR2 pores. The CMRO_2_ response to optogenetic stimulation of pyramidal neurons differed from the CBF response in that it was not augmented by iGluR blockers and was to a great extent reduced by TTX, suggesting dependency upon opening of voltage-gated Na^+^ channels rather than ChR2 pores.

We established a model to explain our optogenetically evoked hemodynamic and metabolic responses based on neuronal connectivity ^16,27^ (fig. 7). During control conditions, optogenetic stimulation directly activated channelrhodopsin-expressing neurons and interneurons and thereby indirectly excited or inhibited their target cells as shown by stimulation-evoked LFPs and gamma activities (fig. 7a, b, top panels). During Optogenetic Pyramidal stimulation, application of iGluR blockers abolished recurrent excitation of neighboring pyramidal neurons and feedback inhibition from neighboring interneurons, as evidenced by the greatly reduced LFP amplitude and evoked gamma activity, respectively (fig. 7b, bottom panel). Addition of TTX to iGluR blockers during Optogenetic Pyramidal stimulation had no further effect on stimulus-evoked LFP, indicating that iGluR blockade had indeed effectively prevented synaptic excitation of pyramidal neurons. The remaining LFP amplitude in the face of iGluR blockade+TTX was likely due to the optogenetically evoked opening of ChR2 pores. We were not able to obtain extracellular LFPs in response to Optogenetic PV stimulation with our experimental set-up. However, we surmise that iGluR blockers prevented excitation of PV interneuron-targeting interneurons ^31^(fig. 7a, middle panel) and that subsequent application of GABA_A_R blockers abolished tonic inhibition and auto-inhibition of PV interneurons (fig. 7a, bottom panel), as suggested by the step-wise augmentations of the CMRO_2_ and CBF responses to Optogenetic PV stimulation in the presence of these blockers.

**Figure 7.**
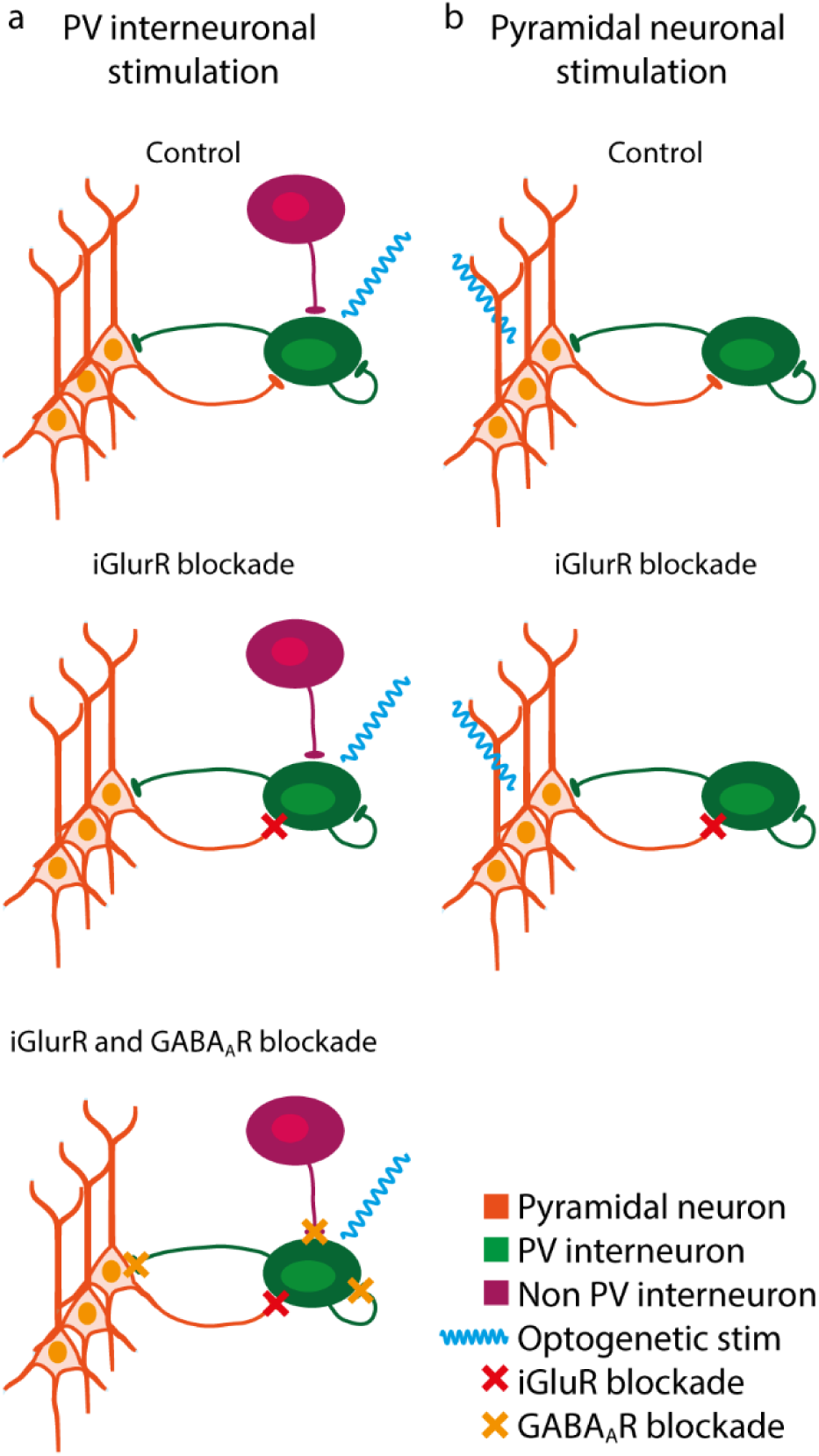
Model of optogenetic stimulation-evoked neuronal and interneuronal activition based on neuronal connectivity patterns in L2/3–L4 of the whisker barrel cortex. Optogenetic PV stimulation shown in (a) and Optogenetic Pyramidal stimulation in (b). In control conditions, Optogenetic PV stimulation (a, upper panel) directly activates PV interneurons (blue wavy line), which in turn inhibit the sparse spontaneous activity of neighboring pyramidal neurons. Combined PV stimulation activates pyramidal neurons as well as PV and non-PV interneurons (not shown). During iGluR blockade (a, middle panel), only PV interneurons are activated both during Optogenetic PV and Combined PV stimulations, as iGluR blockade during Combined PV stimulation prevents recurrent activation of pyramidal neurons as well as activation of their targeted interneurons (not shown). Subsequently, GABA_A_R and iGluR blockade also blocks all inhibitory input via autoreceptors as well as from other PV and non-PV interneurons, potentiating PV interneuron activation during both Optogenetic PV and Combined PV stimulations (a, lower panel). In control conditions, Optogenetic Pyramidal stimulation activates pyramidal neurons directly, bypassing thalamocortical input (b, upper panel). Activation of pyramidal neurons activates PV interneurons, which in turn inhibit pyramidal neurons via recurrent feedback inhibition. This interaction is the basis of gamma activity. During iGluR blockade (b, lower panel), the recurrent feedback inhibitory loop is disrupted, preventing activation of PV interneurons and thereby inhibitory input to pyramidal neurons. Total activation of the system is likely not altered, as the lack of PV interneuron activation may be compensated by a corresponding increase in pyramidal neuron activation.

A prevailing notion concerning brain metabolism is that co-excitation and -inhibition of the same neuron augment cell energy use due to increased flux of both positively and negatively charged ions across the cell membrane; this increase in ion flux may occur with or without affecting neuronal output or transmembrane potential ^17^. In the present study, Combined PV stimulation in which pyramidal neurons simultaneously receive excitatory input from whisker pad stimulation and inhibitory input from activated PV interneurons is therefore expected to allow greater numbers of ions to cross the cell membrane compared to the sum of ions crossing the cell membrane during separate whisker pad stimulation and Optogenetic PV stimulation. CMRO_2_ is thought to represent the work load necessary to restore ion gradients after depolarization ^13,14,32^. In cells entirely dependent upon oxidative phosphorylation to supply energy demands, CMRO_2_ should thus reflect total transmembrane ion flux. In control conditions, we found that the CMRO_2_ response evoked by Combined PV stimulation which activated both pyramidal neurons and PV interneurons amounted to less than the sum of CMRO_2_ responses evoked by separate WP and Optogenetic PV stimulations. In contrast, LFPs evoked by Combined PV stimulation were half as large as those evoked by WP stimulation alone, indicating that net pyramidal neuron excitation was halved by inhibitory input from optogenetically activated PV interneurons. In spite of the increased inhibitory input, the magnitudes of the CMRO_2_ responses to Combined PV stimulation and to whisker pad stimulation were similar, at odds with either the notion that CMRO_2_ reflects total ion flux or that concurrent excitation and inhibition enhance total transmembrane ion flux.

Another conundrum was found during optogenetic stimulation of pyramidal neurons, where CMRO_2_ responses were not augmented by iGluR blockade, but CBF responses were, resulting in a mismatch between CMRO_2_ and CBF. As discussed above, in control conditions, the optogenetically evoked CBF response results from combined reciprocal excitation of pyramidal neurons and recurrent inhibition resulting from collateral activation of all interneuron types, while during iGluR blockade, the evoked CBF response results from excitation of pyramidal neurons alone. LFP amplitude representing net transmembrane ion flux in pyramidal neurons was more than halved by iGluR blockade. The reduction of LFP amplitude here did not indicate that pyramidal neurons received more inhibitory input, as recurrent inhibition was abolished by iGluR blockers. On the contrary, the reduction of LFP amplitude indicated that pyramidal neurons received less excitatory input, due to the lack of reciprocal excitation from neighboring pyramidal neurons. The reduction in excitatory input to pyramidal neurons during iGluR blockade compared to control conditions would be expected to generate a diminished CMRO_2_ response. However, CMRO_2_ was not altered by iGluR blockade despite the putative differences in total transmembrane ion flux. To summarize, Optogenetic Pyramidal stimulation during iGluR blockade evoked less inhibition as well as less excitation compared to control conditions. As described above, in control conditions Combined PV stimulation evoked increased inhibition compared to whisker pad stimulation. Nonetheless, CMRO_2_ remained constant during both procedures. These findings suggest one of the following: 1) that ATP for re-establishing ion gradients is not only supplied by oxidative phosphorylation, but also by aerobic glycolysis ^33^; or 2) that altering the excitatory/ inhibitory current balance affects the per ion energy expenditure for re-establishing ion gradients ^34^; or 3) that the numbers of translocated ions evoked by pyramidal neuron activation are not altered by varying inhibition levels ^17^. In all cases, it appears that CMRO_2_ cannot be used as an a priori indicator of total transmembrane ionic flux.

Substantial gamma activity was only found when either somatosensory or Optogenetic Pyramidal neuron stimulation was included in the stimulation procedure, demonstrating the necessity of pyramidal neuron activation for the induction of gamma activity *in vivo*. Indeed, optogenetic stimulation of PV interneurons alone did not induce gamma activity, but did inhibit the little spontaneous gamma activity that was present in the anesthesized state. Thus, in Pyramidal/ChR2 mice, both somatosensory and optogenetic activation of pyramidal neurons induced proportional gamma activity and CMRO_2_ responses, in line with previous studies showing that gamma oscillations modulate oxygenation of brain tissue ^10,11^. In keeping with previous studies, we found that NMDAR blockade reduced gamma activity and CMRO_2_ responses commensurably in these mice ^35^ and that AMPAR blockade disrupted the relationship between gamma activity and CMRO_2_ (J. A. Cardin et al., 2009), greatly reducing gamma activity evoked by optogenetic stimulation of pyramidal neurons without affecting CMRO_2_. The dissociation between gamma activity and CMRO_2_ was likely due to AMPAR blockade preventing recurrent feedback inhibition as this interaction between pyramidal neurons and PV interneurons is thought to be the basis of gamma oscillations ^1,2,7^. Dissociated gamma activity and CMRO_2_ was also seen in PV/ChR2 mice, where robust CMRO_2_ and gamma responses were evoked in control conditions by WP and Combined PV stimulations. Both stimulation types induced the same CMRO_2_ responses, but decreased gamma activity was found during Combined PV stimulation, where increased inhibition levels were evoked by optogenetic activation of PV interneurons. Thus, gamma activity was reduced both by blocking AMPARs and by increasing inhibitory input to pyramidal neurons, in both cases without affecting CMRO_2_. Keeping in mind the caveats described above, these findings suggest that gamma activity *per se* is not energy consuming, but that the activities of the neuronal components comprising the gamma-inducing circuit are ^8–11^.

In conclusion, we have found that Optogenetic PV stimulation *per se* was able to induce both CMRO_2_ and CBF responses independently of pyramidal neuron activation in good agreement with interneuron participation in neurovascular and -metabolic responses ^36^. The CMRO_2_ and CBF responses to Optogenetic PV interneuronal activation were heavily dependent on the opening of voltage-gated Na^+^ channels. Our data did not support the notion that CMRO_2_ responses to excitation were potentiated by concurrent inhibition. Lastly, we found that pyramidal neuron activity was necessary for the generation of gamma activity *in vivo* and that oxygen consumption is largely driven by neuronal activation and not by the synchronized interplay between neuron types, in the form of gamma activity, as has previously been suggested.

## Material and Methods

### Animal handling

All procedures involving animals were approved by the Danish National Committee according to the guidelines set forth in the European Counciĺs Convention for the Protection of Vertebrate Animals used for Experimental and Other Scientific Purposes. Mutant knock-in transgenic mice expressing channel rhodopsin 2 (ChR2) ^37^ in either parvalbumin positive interneurons (PV/ChR2 mice, *n*=9) or in Ca^2+^/calmodulin-dependent protein kinase IIα (CamK2α) positive pyramidal neurons (PYR/ChR2 mice, *n*=9) were used. These mice were generated by cross-breeding B6.Cg-*Gt(ROSA)26Sor^tm32(CAG–COP4*H134R/EYFP)Hze^*/J with either B6.129P2-*Pvalb^tm1(cre)Arb^*/J or with B6.Cg-Tg(Camk2a-cre)T29-1Stl/J mice strains. All three strains (Jackson Laboratory, USA) were homozygous and F1 progeny, mice of both sexes, 4-6 mo old were used.

Animals were anaesthetized with xylazine + ketamine (induction:10 + 60 mg/kg i.p.). During surgery, supplementary doses of ketamine (30 mg/kg i.p.) were given every 20 min. The trachea was cannulated for mechanical ventilation (SAR-830/P, CWE Inc., Pennsylvania, USA) and catheters were placed in the left femoral vein and artery for measuring arterial blood pressure and blood gases and for infusion of anaesthesia. A metal plate with a hole (5 mm in diameter) was glued to the scull with cyanoacrylate glue (Loctite®, Henkel, Germany) with accelator (Insta-Set®, Bob Smith Industries, CA, USA). A 4-mm-diameter craniotomy was drilled over the sensory barrel cortex (0.5 mm behind and 3 mm to the right of bregma). The dura was removed and the brain was covered with artificial cerebrospinal fluid (aCSF; in mM, NaCl 120, KCL 2.8, NaHCO_3_ 22, CaCl_2_ 1.45, NaHPO_4_ 1, MgCl_2_ 0.876, glucose 2.55, buffered with HEPES (2.38 g/l); pH = 7.4 at 22 °C).

After surgery, anaesthesia was switched to α-chloralose (α-chloralose-HBC complex, 0.5g/ml, 0.01 ml/10 g/h i.v.). At the end of the experimental protocol, the mouse was immediately euthanized with ketamine followed by decapitation.

To ensure stable physiological conditions of the mice during the experiment, blood pressure (pressure monitor BP-1; World Precision Instruments) and end expiratory CO_2_ (Capnograph type 340; Harvard Apparatus) were monitored continuously and two arterial blood samples were drawn to assess blood gases (pO_2_, 95–140 mmHg; pCO_2_, 30–45 mmHg; pH, 7.3–7.45, ABL 700Series radiometer). Body temperature was maintained at 37°C using a rectal temperature probe and a heating blanket (TC-1000 Temperature Controller; CWE).

### Stimulation protocol

All stimulations were controlled by a sequencer file running within the Spike2 software (version 7.2, CED, Cambridge, England) and the stimulation period was 15 s.

### Whisker-Pad stimulation (WP stim)

The mouse whisker barrel cortex was activated by stimulating the contralateral infraorbital branch of the trigeminal nerve (1.5 mA, 2 Hz, pulse length of 1 ms). A set of custom-made bipolar electrodes was inserted percutaneously into the infraorbital foramen (cathode) and the ipsilateral masticatory muscles (anode). Stimulation of the infraorbital nerve corresponds to stimulation of the entire ipsilateral whisker pad (fig 1e).

### Optogenetic stimulation of PV interneurons or pyramidal neuron in the whisker barrel cortex (Optogenetic PV/Pyramidal stimulation)

A laser fibre transmitting blue laser light (473 nm, 2 mW by the fibre tip; DL 473-050-O, CrystaLaser, NV, USA) was placed just above the whisker barrel cortex. The stimulation parameter for Pyramidal/ChR2 mice were 2 Hz with a pulse length of 100 ms and for PV/ChR2 were they 100 Hz, 7,5 ms (fig 1c, f).

### Combined WP and optogenetic stimulation (Combined PV/Pyramidal stimulation)

In Pyramidal/ChR2 mice using combined sensory and optogenetic stimulation protocols, stimulation pulses occurred simultaneously. In PV/ChR2 mice using combined protocols, stimulation pulses were in phase but occurred simultaneously once out of every 50 pulses due to different stimulation frequencies (fig 1d, g).

### Experimental protocol

In PV/ChR2 mice, WP, Optogenetic PV and Combined PV stimulations were performed during control conditions and in the presence of the NMDAR blocker MK-801 (300 µM) + the AMPAR blocker NBQX (200µM), followed by the addition of the GABA_A_R blocker, GABAzine (20 µM), and lastly, by the addition of the voltage-sensitive Na^+^ channel blocker TTX (20 µM). In Pyramidal/ChR2 mice, WP, Optogenetic Pyramidal and Combined Pyramidal stimulations were given during control conditions and in the presence of MK-801 alone, followed by the addition of NBQX, and lastly by the addition of TTX.

All blockers were applied topically to the cortex 20 minutes before stimulation was commenced. For each combination of blockers, 3 WP stimulation trains, 3 combined stimulation trains and 3 optogenetic stimulation trains were applied in random order. Each stimulation train of 15 s was preceded by a 30 s baseline and followed by a 120 s recovery time (fig 1h).

TTX was obtained from VWR International (Søborg, Denmark), all other substances were obtained from Sigma-Aldrich (Copenhagen, Denmark).

### Electrophysiology

The total electrical signal (0.5–3000 Hz) was recorded with a vertical 16-channel linear multiarray probe with 50 µm between electrode sites (NeuroNexus, Michigan, USA) and a 16-channel amplifier (gain x1000, bandwidth 1-10 000 Hz; PGA16, Multichannel system). The probe was inserted vertically into the barrel cortex with the top electrode at the level of the pial surface and the bottom electrode extending to a depth of 750 µm, encompassing the barrel cortex from layers 1 to 5. Electrical analogue signals were digitally sampled at least five times the low-pass filter frequency using a Power1401 mk II interface (CED, Cambridge, UK).

#### Extracellular local field potentials (LFP)

To obtain LFPs, the total electrical signal was low-pass filtered at 300 Hz. The LFP for each stimulation train was averaged across stimulation impulses and LFP amplitude was taken as the greatest negative deflection occurring within 20 ms after the stimulus artefact. LFP amplitudes were then averaged across stimulation trains for each stimulation modality and treatment.

#### Gamma oscillations

To calculate the power of gamma oscillatory activity (30-90 Hz; gamma power), stimulation artefacts were removed from the total electrical signal. A Fourier transformation in Matlab (function *bandpower*) calculated the gamma power for each stimulation train and preceding 15 s of baseline in layer 2/3 corresponding to the cortical depth of 250 µm. Mean baseline and stimulation-evoked gamma powers were averaged across stimulation trains, then across mice, for each stimulation modality and treatment.

### Laser speckle contrast imaging

Laser speckle contrast imaging (LSCI) was performed using infrared coherent light (785 nm, LP 785- SF 100, Thorlabs) controlled by a diode driver (CLD 1011, Thorlabs). Laser diode output power was 40 mW and illuminated area was approximately 4 cm^2^, resulting in the power density of approx 10 mW/cm^2^ at the surface of the cortex ^38^. A CMOS camera (acA2000-165umNIR, Basler, Germany) equipped with a zoom lens (VZMTM 450i Zoom Imaging Lens, Edmund Optics, Germany) was used for signal collection. Exposure time was set to 4 ms and field of view to 800 x 800 pixels. Zoom was 1x leading to digital resolution of 5 μm per pixel. To prevent contamination of the laser speckle signal by laser light emitted during optogenetic stimulation, the lens was fitted with a red color filter with a cut-on wavelength of 600 nm (#46-539, Edmund Optics, Germany).

Raw data images were acquired at 25 fps. To estimate the so-called “blood flow index”, temporal laser speckle contrast analysis was applied ^39^, resulting in the reduction of frame rate to 1 frame per second while preserving the spatial resolution of the raw data. Blood flow dynamics were averaged over a circular roi (Ø 40 pixels) corresponding to the cortical area monitored by the multiarray probe and oxygen electrode.

### Local tissue oxygen partial pressure (tpO_2_)

tpO_2_ was measured amperometrically in layer 2/3 at cortical depth of 250 μm using a modified Clark-type polarographic oxygen electrode (OX-10, tip diameter: 10 μm, field of sensitivity diameter: 20 μm; Unisense A/S, Denmark). The oxygen electrode was calibrated in air-saturated and oxygen-free saline (0.9%) at room temperature. The oxygen electrode was connected to a high-impedance picoamperemeter (PA 2000, Unisense A/S), and oxygen signals were A/D converted, sampled at 100 Hz using a Power 1401 A/D converter (CED) running with Spike2 software (CED), and averaged to give an effective sampling rate of 1 Hz.

#### Calculation of CMRO_2_

CMRO_2_ was calculated from tpO_2_ and CBF ^40,41^. The relationship between these three variables is given by:

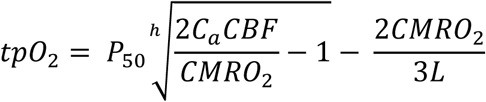

where *P_50_* is the half-saturation tension of the oxygen-hemoglobin dissociation curve, *h* is the Hill coefficient of the same dissociation curve, *C_a_* is the arterial oxygen concentration and *L* is the effective diffusion coefficient of oxygen in brain tissue. The value of *L* was determined from baseline tpO_2_ values of mice included in the present study and from baseline CBF and CMRO_2_ values of mice in similar conditions of anesthesia, which in the literature were reported to be 73 ml·100 g^−1^·min^−1^ and 263 μmol·100 g^−1^·min^−1^ ^42^. *L* was calculated to be 5.39 μmol·100 g^−1^·min^−1^·mmHg^−1^ for standard values of *P*_50_ (41 mmHg, ^43^, *h* (2.7), and *C_a_* (8 μmol ml^−1^).

### Immunohistochemistry

Immunohistochemistry was used to evaluate the co-expression of yellow fluorescent protein (GFP)- tagged ChR2 with PV, respectively CaMKIIα, in the whisker barrel cortex. Mice deeply anesthetized with ketamine+xylazine (0.15 mg+0.01 mg/g body weight) were perfused with 4% paraformaldehyde (PFA). The brains were post-fixed overnight in 4% PFA, transferred to 30% sucrose in 0.1 M phosphate buffered saline (PBS) for 2 – 3 days and sectioned (20 μm) with a cryostat. Free floating sections from PV/ChR2 mice were blocked with 0.3% Triton X-100 and 3% normal goat serum (NGS) in PBS (blocking solution PV) for 1 hour at RT, incubated overnight with chicken anti-GFP antibody (AB13970, Abcam, 1:600) and rabbit anti-parvalbumin antibody (AB11427, Abcam, 1:1000) at 4 °C followed by incubation the next day with goat anti-chicken antibody (647nm, A21449, ThermoFisher, 1:600) and goat anti-rabbit antibody (568 nm, A11036, ThermoFisher, 1:1000). Free floating sections from ChR2/CaMKIIα mice were blocked with 0.2% Triton X-100, 5% NGS and 2.5% bovine serum albumin in PBS (blocking solution CaMKIIα) for 1 hour at RT. A sequential protocol was employed, where the sections were incubated 36 hours with rabbit anti-CaMKIIα antibody (SAB4503250, Sigma-Aldrich, 1:50) at 4 °C, extensively washed and then incubated with goat anti-rabbit antibody (as above) at RT for 2 hours. After this, the sections were incubated with primary chicken anti-GFP antibody (as above) overnight at 4 °C followed by incubation with goat anti-chicken antibody (as above) at RT for 2 hours. Hoechst (H3570, Invitrogen, 1:5000) was added for 7.5 minutes during one of the washes. All antibodies were dissolved in blocking solution of the mouse strain being evaluated. Slices were mounted on glass slides (Superfrost Plus Gold, Menzel-Gläser, Thermo Scientifica) with Slowfade Diamond Antifade Mountant (Invitrogen). Images were captured using an upright laser scanning confocal microscope (LSM 700 or 710, Zeiss) at 20x or 63x magnification.

### Calculations and Statistics

Despite the cut-off filter preventing contamination of the laser speckle signal by light with wave lengths under 600 nm, we found that optogenetic stimulation of ChR2 triggered fluorescence of the fused EYFP protein and interfered with recordings from PYR/ChR2 mice, but not PV/ChR2 mice. Accordingly, no data points for PYR/ChR2 mice are shown during optogenetic stimulation in CBF and CMRO_2_ time lines. However, when quantifying time line curves for PYR/ChR2 mice, areas under the curve (AUCs) included estimations of the responses during stimulation achieved by linearly interpolating the last baseline points with the first points after the termination of the optogenetic stimulation. This approach was adopted to be able to compare responses between stimulation modalities and between mouse strains.

CBF and CMRO_2_ is given as percent of baseline (% baseline) in time line plots and as % baseline*s in AUC graphs. AUC was calculated as the integral of the positive CBF or CMRO_2_ response from start of stimulation to return to baseline (fig. 1i, j).

Linear mixed effects modeling (MEM) analyses were performed in R using ^44^ functions *lme4* ^45^ or *glmmPQL* ^46^. As the data comprised measurements from many time points from each animal, MEM was employed to account for the non-independence of the data. Treatment, stimulation type, and their interaction were included as fixed effects; mouse was included as random effect. Bonferoni corrections were applied to post hoc *t*-tests. α was taken as 0.05. Data are shown as mean ± standard error of the mean.

## Acknowledgements

The authors wish to thank Thomas Hartig Braunstein, PhD and Pablo Hernandez-Varas, PhD at the Center for Integrative Microscopy, University of Copenhagen for their great help with IHC imaging and image analyses; and Micael Lønstrup for his invaluable help with all technical support in the laboratory. This study was supported by the Center for Healthy Aging, University of Copenhagen; the Lundbeck Foundation; the Novo-Nordisk Foundation; and the Danish Council for Independent Research.

