## Supplementary figures for "Modification of oxygen consumption and blood flow in mouse somatosensory cortex by cell-type-specific neuronal activity"

**S1**

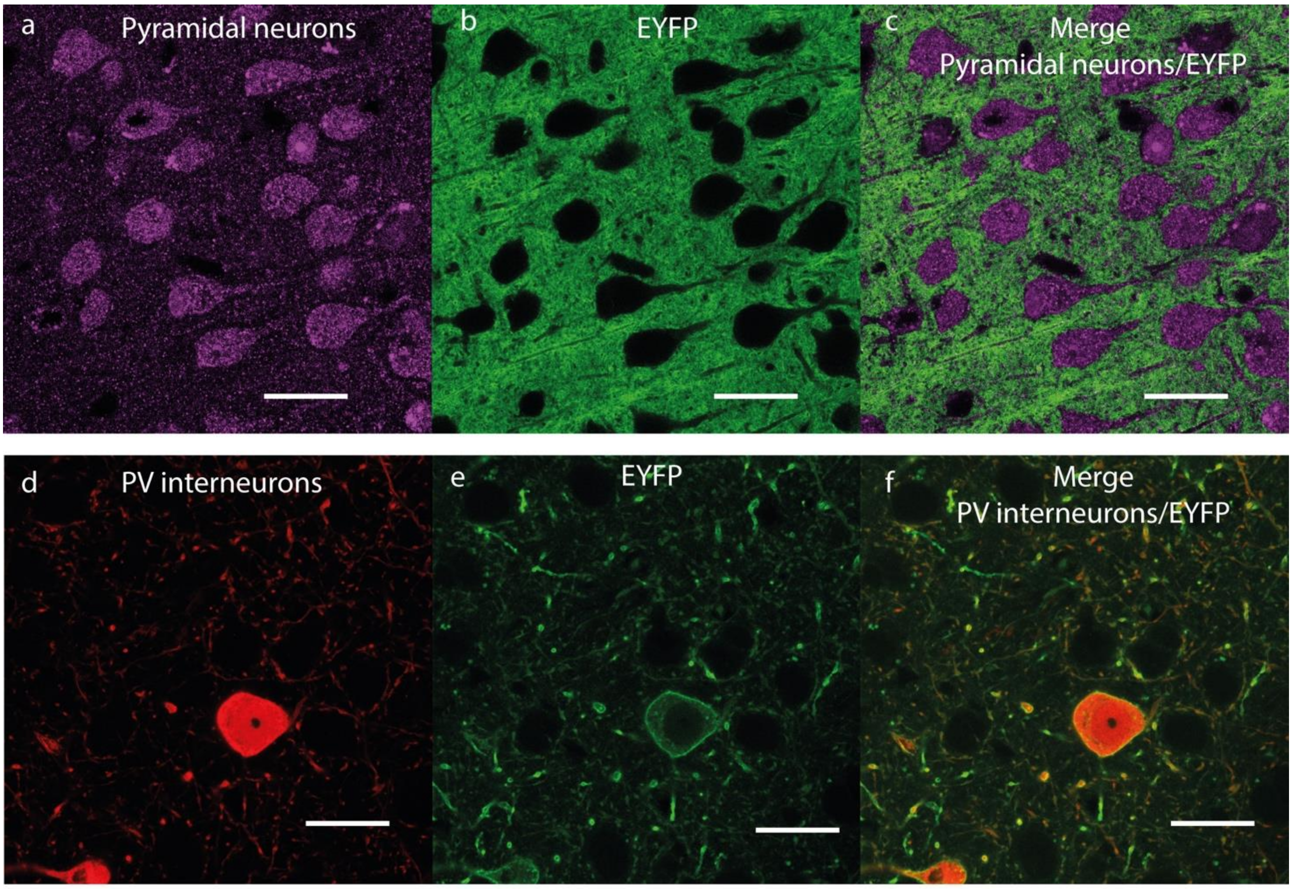


**S1. IHC images of the barrel cortex of Pyramidal/ChR2 and PV/ChR2 mice.** Pyramidal/ChR2 WB cortex L2/3 shown in panels (a – c) and PV/ChR2 WB cortex L2/3 shown in panels (d – f). (a) CamKIIα labelling of pyramidal neurons, (d) parvalbumin labelling of PV interneurons, (b, e) labelling of EYFP tethered to ChR2, (c) merged CamKIIα and EYFP-ChR2 images from (a, b), (f) merged PV and EYFP-ChR2 images from (d, e). Note the massive labelling of EYFP-ChR2 in CamKIIα neuronal processes compared to the limited labelling of EYFP-ChR2 in PV interneuronal processes (b, e).

**S2**

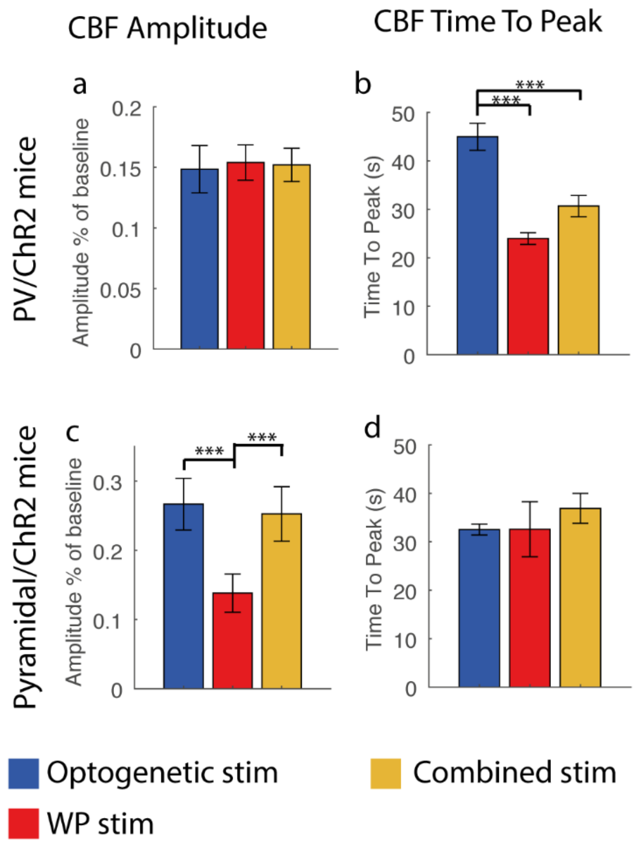


**S2: Maximal amplitude and time-to-peak for CBF time courses during control conditions.** Maximal amplitude for (a) PV/ChR2 and (c) PYR/ChR2 mice. Time-to-peak for (b) PV/ChR2 mice and (d) PYR/ChR2 mice. (a) During control conditions, maximal amplitudes of CBF time courses were similar between stimulation types in PV/ChR2 mice. (b) Time-to-peak during control conditions did differ, being delayed by ~21 s during Optogenetic PV stimulation compared to WP stimulation, suggesting a poly-synaptic or non-ionotropic signalling pathway for optogenetic stimulation-evoked CBF responses in PV/ChR2 mice. (c) While the maximal amplitude of the CBF time course was greatly reduced during WP stimulation of pyramidal cells in PYR/ChR2 mice compared to optogenetic and combined Pyr stimulations of the same pyramidal neurons, (d) time-to-peak did not differ.

Data is given by means ± SEM. Significant differences are shown by asterisk(s): *P < 0.05, **P < 0.01 and ***P < 0.005.

**S3**

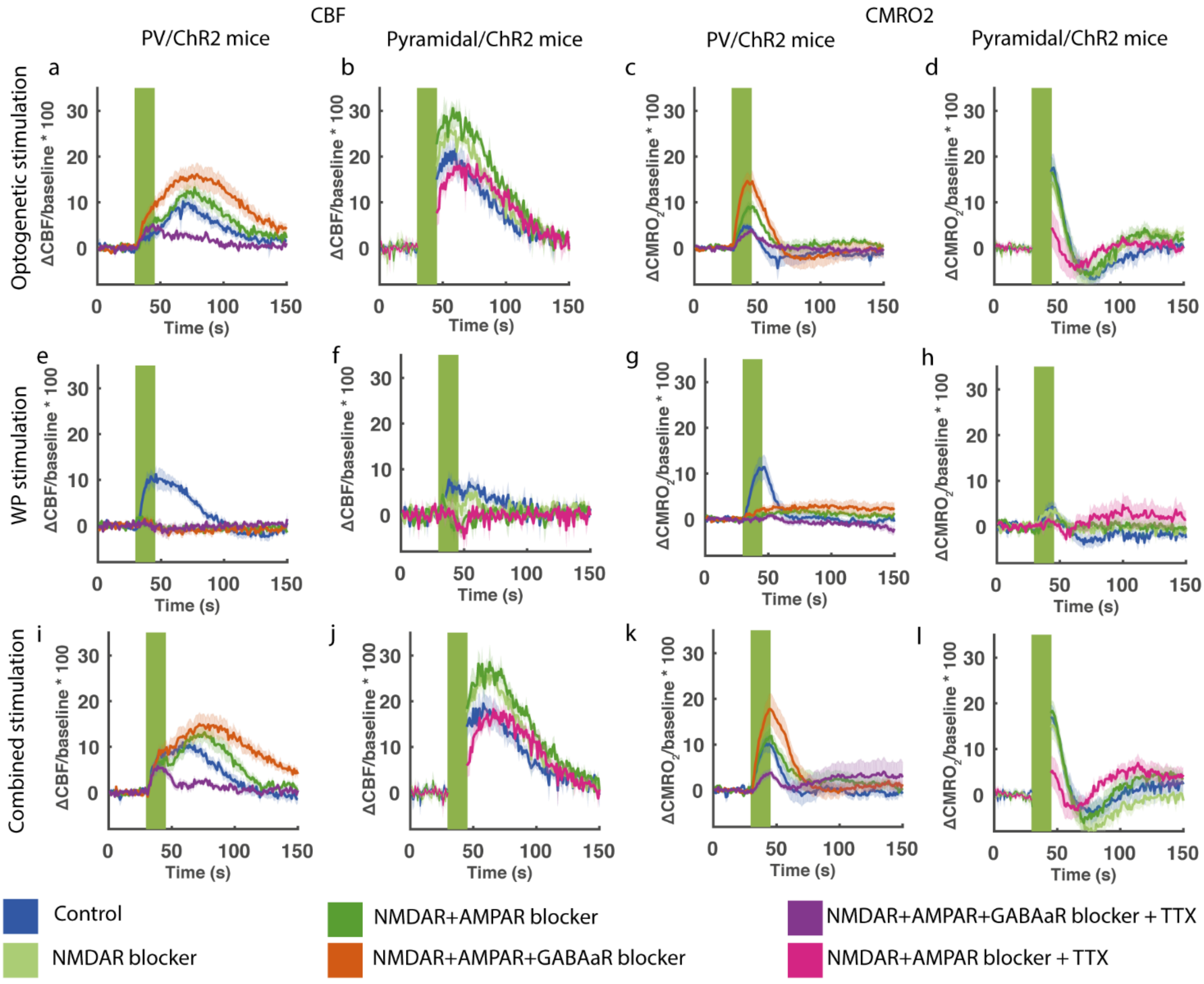


**S3: Time course for the different mice strains and treatments.** Optogenetic stimulation-evoked CBF and $CMRO_2$ responses in (a, c) PV/ChR2 mice and (b, d) PYR/ChR2 mice. WP stimulation-evoked CBF and $CMRO_2$ responses in (e, g) PV/ChR2 mice and (f, h) PYR/ChR2 mice. Combined stimulation-evoked CBF and $CMRO_2$ responses in (i, k) PV/ChR2 mice and (j, l) PYR/ChR2 mice. (a-l) Graphs show response time courses. During optogenetic and combined stimulations (a, c, i, k), treatment with NMDAR+AMPAR+$GABA_aR$ blockers in PV/ChR2 mice augmented both CBF and $CMRO_2$ responses. TTX abolished the second part of the biphasic Optogenetic PV stimulation-evoked CBF time course, leaving the first part unaffected (a, i). In contrast to PV/ChR2 mice, NMDAR+AMPAR blockers augmented CBF responses to Optogenetic and Combined Pyramidal stimulation in PYR/ChR2 mice but had no effect on $CMRO_2$ responses and TTX did not abolish CBF responses (b, d, j, l). The iGluR blockers abolished CBF and $CMRO_2$ responses to WP stimulation (e,f,g,h). The CBF and $CMRO_2$ stimulation time courses are averaged across animals. Each CBF time course was extracted from a roi (40 pixels in diameter) placed next to the insertion points of the oxygen electrode and the multichannel electrode in the laser speckle recordings and used for calculating $CMRO_2$ time courses, $AUC_{CBF}$ and $AUC_{CMRO2}$. Green fields represent 15 s stimulation trains. All data is presented as mean±SEM. n = 9 for all groups. Significant difference is shown by asterisk(s): *P < 0.05, **P < 0.01 and ***P < 0.005.
